# Parallel sexual and parasexual population genomic structure in *Trypanosoma cruzi*

**DOI:** 10.1101/338277

**Authors:** Philipp Schwabl, Hideo Imamura, Frederik Van den Broeck, Jaime A. Costales, Jalil Maiguashca, Michael A. Miles, Bjorn Andersson, Mario J. Grijalva, Martin S. Llewellyn

## Abstract

Genetic exchange and hybridization in parasitic organisms is fundamental to the exploitation of new hosts and host populations. Variable mating frequency often coincides with strong metapopulation structure, where patchy selection or demography may favor different reproductive modes. Evidence for genetic exchange in *Trypanosoma cruzi* over the last 30 years has been limited and inconclusive. The reproductive modes of other medically important trypanosomatids are better established, although little is known about their variability on a spatio-temporal scale. Targeting a contemporary focus of *T. cruzi* transmission in southern Ecuador, we present compelling evidence from 45 sequenced genomes that *T. cruzi* (discrete typing unit I) maintains sexual populations alongside others that represent clonal bursts of parasexual origin. Strains from one site exhibit genome-wide Hardy-Weinberg equilibrium and intra-chromosomal linkage decay consistent with meiotic reproduction. Strains collected from adjacent areas (>6 km) show excess heterozygosity, near-identical haplo-segments, common mitochondrial sequences and levels of aneuploidy incompatible with Mendelian sex. Certain individuals exhibit trisomy in as many as fifteen chromosomes. Others present fewer, yet shared, aneuploidies reminiscent of mitotic genome erosion and parasexual genetic exchange. Genomic and intra-genomic phylogenetics as well as haplotype co-ancestry analyses indicate a clear break in gene-flow between these distinct populations, despite the fact that they occasionally co-occur in vectors and hosts. We propose biological explanations for the fine-scale disconnectivity we observe and discuss the epidemiological consequences of flexible reproductive modes and their genomic architecture for this medically important parasite.

## INTRODUCTION

*Trypanosoma cruzi* is a kinetoplastid parasite and the causative agent of zoonotic Chagas disease (CD) in Latin America, where ca. six million people are currently infected (1). Mucosal or abrasion contact with the infected feces of hematophagous triatomines constitutes the primary mode of *T. cruzi* transmission (2). Infection with *T. cruzi* results in chronic CD in 30 - 40% of cases, characterized by a spectrum of fatal cardiac and intestinal pathologies. Early-stage acute CD can also be fatal, especially among infants and in orally transmitted outbreaks of the disease (3, 4). *T. cruzi* transmission is a zoonosis maintained via complex transmission cycles involving numerous species of triatomine insects (5) and hundreds of different species of mammals (6). Transmission cycles can thus differ markedly across the parasite’s wide distribution in the Americas at macro and micro scales.

The Trypanosomatidae, which include several trypanosomatids of medical and veterinary importance – e.g., *Trypanosoma brucei* ssp., *Leishmania* spp., *Trypanosoma vivax* and *Trypanosoma congolense*, in addition to *T. cruzi* – are a monophyletic group of obligate parasites (7). The Trypanosomatidae are basal eukaryotes and share many biological idiosyncrasies, including the kinetoplast (8) and polycistronic transcription control (9). Despite their basal status, the Trypanosomatidae share much of the core meiotic machinery of ‘higher’ eukaryotes (10). Nonetheless, evidence of the extent to which such machinery might remain active in trypanosomatids has been difficult to obtain and often conflicting despite the importance of such process in the spread of epidemiologically relevant traits (e.g., drug resistance or virulence in different hosts (11–13)). Frequent meiotic recombination in *T. b. brucei* has been established in the laboratory and field for several decades (14–16). More recently, genome-scale signatures of meiosis have also been detected in *T. congolense* (17). In contrast, robust genomic evidence suggests the closely related *T. b. gambiense* is completely asexual (18). Life histories in *Leishmania* seem no less complex, with alternating sexual and asexual recombination proposed to explain population genomic structures (19), although meiotic exchange is well supported in experimental crosses (20).

*T. cruzi* is the last of the ‘Tritryps’ for which the extent and mechanism of genetic exchange remains to be fully elucidated. Limited evidence for genetic exchange has been observed in the field (21), although inappropriate study designs and markers systems of insufficient resolution hamper interpretation of the data (11). Furthermore, the parasexual mechanism of genetic exchange proposed in the laboratory – one of genotypic ‘fusion-then-loss’ (22) – has been difficult to reconcile with field research via measurement of somy variation in contemporary populations or genetic composition of ancient hybrid lineages (21, 23–26). This lack of clarity has lead some to propose *T. cruzi* as a paradigm for ‘predominant clonal evolution’ in parasitic protozoa (27, 28) – an idea which may not reflect biological reality.

To address this major knowledge gap in the biology of trypanosomatids, we generated whole-genome sequence data from 45 *T. cruzi* Discrete Typing Unit I clones, as well as several non-cloned *T. cruzi* strains, collected from triatomine vectors and mammalian hosts in an endemic transmission focus in Loja Province, southern Ecuador. After mapping sequences against a recent PacBio sequence assembly (29), we explored patterns of population structure and genetic recombination. Our data reveal that *T. cruzi* does indeed reproduce sexually at high frequency via a mechanism consistent with classical meiosis. However, we also demonstrate that parasite populations with radically distinct, parasexual population structures can exist in near-sympatry alongside those that are panmictic. Our data make a significant contribution towards the consolidation of current theories around genetic exchange in the Trypanosomatidae.

## RESULTS

### Extensive divergence between sympatric parasites

Paired-end sequence reads from 45 single-strain and 14 non-cloned *T. cruzi* cultures aligned to the *T. cruzi* TcI X10/1 Sylvio genome assembly at 27x average mapping depth. After variant filtration and masking (see *Materials and Methods*), 206,619 SNP sites were identified against the reference. Excluding non-cloned *T. cruzi* cultures, 130,996 diagnostic markers were identified that clearly separated the 45 clones into two phylogenetic clusters (Fig. 1a and Fig. S1a). Cluster 1 contained 15 of 17 isolates from the ‘Bella Maria’ (BM) sampling region. Cluster 2 contained all isolates from nearby ‘Ardanza’ (AR; n = 11; <7 km from BM), as well as isolates from ‘El Huayco’ (EH; n = 12) and ‘Gerinoma’ (GE; n = 3) study sites ca. 35 km northwest of BM (Fig. 1a (inset); see also spatial coordinates in Tbl. S1). Unsupervised k-means cluster selection (30) confirmed a single major partition (i.e., k = 2) in the data, though mild improvements to model fit continued through k = 6 (Fig. S8a). To describe such additional substructure, we reconstructed each phased genome as a mosaic of haplo-segments sharing ancestry with other samples of the dataset (including non-cloned samples) (31). In the resultant co-ancestry matrix (plotted as a heatmap with *F_ST_* scores in Fig. 1b), intensity of haplotype-sharing increased within both clusters relative to the spatial origin of each isolate, with the exception of TCQ_3087 (sampled in BM but associated to Cluster 2) and TRT_3949 clones (sampled in Santa Rita (SR), near EH, but associated to Cluster 1). Four non-cloned samples TBM_4307_MIX, TBM_3131_MIX, TRT_4082_MIX (cultured from *Rhodnius ecuadoriensis*) and MBC_1529_MIX (cultured from *Simosciurus nebouxii*) showed shared ancestry across clusters, indicating that parasites from divergent Clusters 1 and 2 co-occur in the same vectors and hosts.

**Figure 1.**
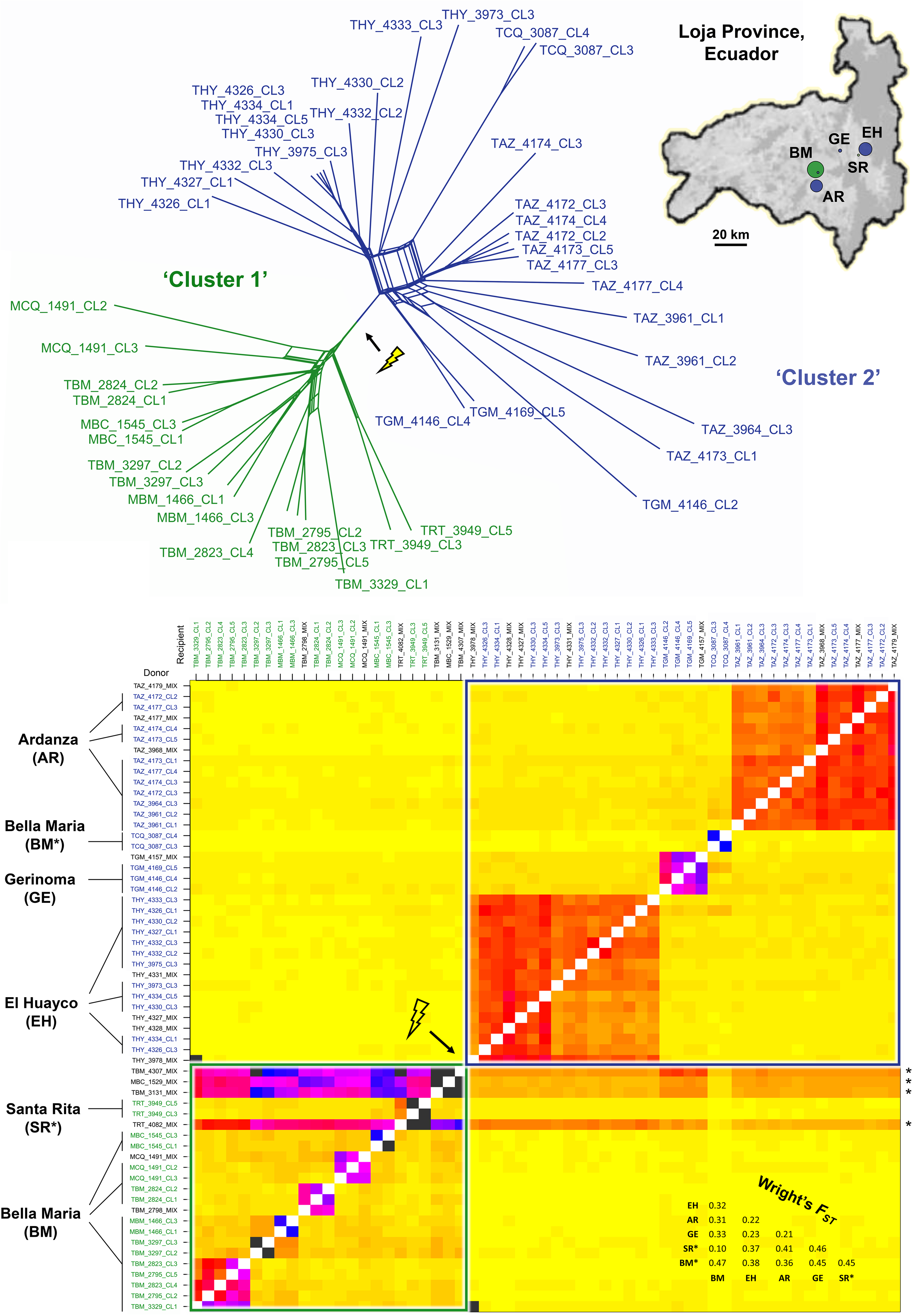
Phylogenomic relationships among *T. cruzi* I isolates from southern Ecuador. (**a**) Data are represented as a split network by the Neighbor-Net algorithm (82). Pairwise genetic distances are defined as the proportion of non-shared genotypes across all SNP sites for which genotypes are called for >40 individuals. Arrow (and flash) indicate a strong, unambiguous break in gene-flow between two reticulate assemblages, ‘Cluster 1’ (green) and ‘Cluster 2’ (blue). (**b**) Heatmap of co-ancestry based on a sorted haplotype co-ancestry matrix *x_ij_*, which estimates the number of discrete segments of genome *i* that are most closely related to the corresponding segment of genome *j*. These ‘nearest-neighbor’ relationships from fineSTRUCTURE (31) analysis are sorted such that samples clustered along the diagonal are those that most share recent genealogical events and pairwise comparisons outside of the diagonal indicate levels of genetic connectivity among these clusters. The matrix also includes ‘genomes’ of non-cloned *T. cruzi* I cultures. Strong horizontal banding points to the accumulation of diversity from throughout the dataset in three of these original infections. The upper right inset shows sampling regions in Loja Province, Ecuador. Region labels are abbreviated as in the co-ancestry matrix. Point sizes correspond to sample sizes and colors correspond to cluster membership (see network). Estimates of Wright’s *F_ST_* (Weir and Cockerham’s method) at various subdivisions are provided on the lower right.

### Sympatric Mendelian and non-Mendelian population genetic signatures

To explore eco-evolutionary processes potentially underpinning sympatric divergence in *T. cruzi*, we established population genetic structure in each partition. In BM isolates of Cluster 1, allele frequencies at variable loci matched those predicted for random mating, with inbreeding coefficients distributed thoroughly over zero (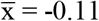, σ = 0.38; Fig. S2) and 87,600 of 96,691 (91%) variant loci meeting expectations for Hardy-Weinberg equilibrium (Tbl. 1). Heterozygosity was unevenly distributed (e.g., Fig. S3), often interrupted by long runs (>100 kb) of homozygosity (4,675 instances in total (Tbl. S2)) and appeared fixed at only 4% (2,134 / 58,102) polymorphic sites (Tbl. 1). Singl-enucleotide variants were often found exclusive to few or single samples, with >50% variants found in <5% individuals and 14% sites comprising singleton SNPs. Phased 30 kb sequence alignment thus yielded a genome-wide average of 10.3 distinct haplotypes across the BM group (for windowed haplotype counts on chromosome 1, see Fig. S4a).

**Table 1.** Population genetic descriptive metrics for *T. cruzi* I isolates from Bella Maria (BM, associated to Cluster 1), El Huayco (EH, Cluster 2) and Ardanza (AR, Cluster 2). Abbreviations: Poly. (Polymorphic); MAF (within-group Minor Allele Frequency); HWE (Hardy-Weinberg Equilibrium); Het. (Heterozygous).

| Group (n) | Poly. Sites | Median Nucleotide Diversity ($\pi$) per site | Median Watterson Estimator ($\theta$) per site | % Poly. Sites at MAF > 0.05 | Private Sites (vs. BM / EH / AR) | Singleton Sites | Poly. Sites in HWE | Het. Sites | Fixed Het. Sites |
| --- | --- | --- | --- | --- | --- | --- | --- | --- | --- |
| BM (15) | 96,691 | 0.09 | 0.001 | 48 % | 0 / 40,177 / 40,262 | 14,013 | 87,500 | 58,102 | 2,134 |
| EH (12) | 80,052 | 0.15 | 0.001 | 70 % | 23,538 / 0 / 18,016 | 4,525 | 33,980 | 58,980 | 44,945 |
| AR (11) | 78,325 | 0.16 | 0.001 | 71 % | 21,896 / 16,289 / 0 | 6,064 | 35,799 | 58,392 | 45,287 |

The extent and pattern of allelic diversity in Cluster 2 was highly distinct to that observed in Cluster 1. In EH and AR, heterozygosity accounted for 73% and 74% total SNP-variation, respectively, leading to Hardy-Weinberg departures at 42% (EH) and 46% (AR) total sites (Tbl. 1). High levels of heterozygosity extended continuously across all chromosomes (e.g., Fig. S3). Long runs of homozygosity affected just 10 of 23 samples (17 instances in total (Tbl. S2)) and heterozygous polymorphism featured largely as non-segregating sites. Seventy-six per cent (44,945 / 58,980) of heterozygous loci occurred as fixed SNPs within EH and 78% (45,287 / 58,392) occurred as such in AR. Rare (i.e., observed in <5% individuals) single-nucleotide variants accounted for only 30% (EH) and 29% (AR) total variation. Accordingly, haplotype diversity remained low, with a genome-wide average of 5.6 distinct haplotypes found in phased 30 kb sequence alignment within both EH and AR (see windowed counts for chromosome 1 in Fig. S4a). Tajima’s *D* values were highly inflated compared to those in BM (Fig. S5).

### Linkage decay and rates of meiotic recombination in *T. cruzi*

In sexually recombining organisms, pairwise SNP-associations (r^2^) are predicted to decay with map distance due to crossover that occurs between chromosomes during meiosis. We plotted r^2^ against pairwise map distance for all diagnostic SNP loci identified at BM. Fig. 2a depicts results for chromosome 1, with linkage declining sharply in the first few kilobases, then more gradually and approaching zero near 60 kb. Linkage decay was apparent on other chromosomes examined (chromosomes 1, 5, 21 and 26 (Fig. 2b), chosen for their reliability in mapping). A similar pattern also emerged in analysis restricted to ‘core’ sequence regions (genes syntenous to *T. b. brucei* and *L*. *major*) (suppl. Fig. S6). In contrast to BM, analyses of linkage decay for EH and AR showed no relationship between r^2^ and map distance. Rather, complete and intermediate linkage, as well as an abundance of random variant-associations, featured continuously through all distance classes on the same chromosomes surveyed in BM (e.g., chromosome 1, Fig. 2c-d).

**Figure 2.**
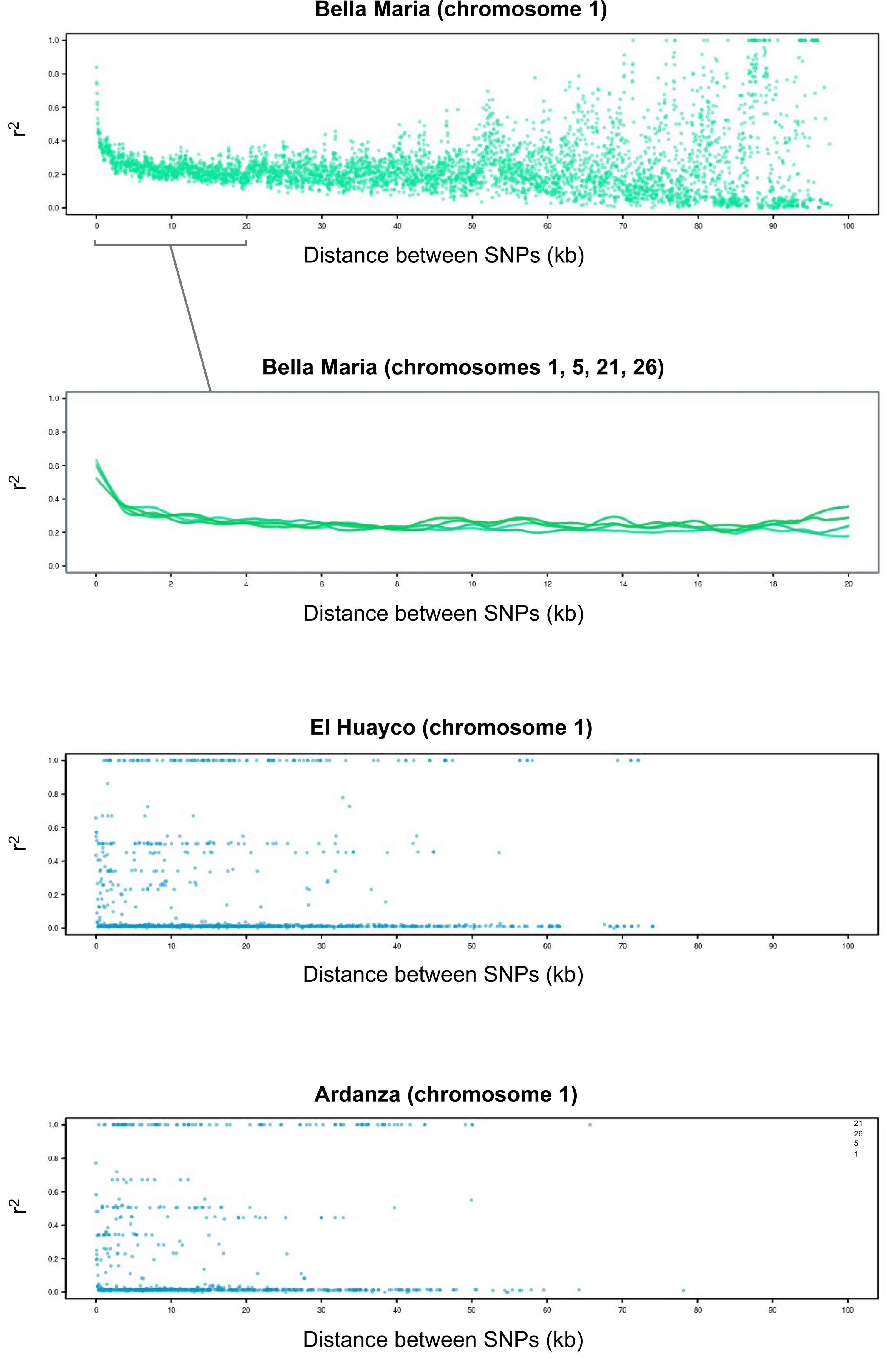
Linkage decay and different rates of recombination in *T. cruzi* I populations. (**a**) Decay of linkage disequilibrium on chromosome 1 for *T. cruzi* I isolates from Bella Maria. Average pairwise linkage values (r^2^) among all diagnostic SNP sites are plotted for map distance classes between 0 and 100 kb. (**b**) Local regression curves for the decay of linkage disequilibrium on chromosomes 1, 5, 21 and 26 for *T. cruzi* I isolates from Bella Maria. **(c-d)** Lack of linkage decay on chromosome 1 for *T. cruzi* I isolates from El Huayco and Ardanza. Average pairwise linkage values (r^2^) are plotted against distance classes as for Bella Maria above.

We estimated the frequency of meiosis (*N_ρ_* / *N_θ_*) in our study populations by comparing two different estimates of effective population size. The first estimate, *N_ρ_*, is based on recombinational diversity observed in the sample and represents the number of cells derived from mating. The second, *N_θ_*, is based on mutational diversity and represents the total number of cells, irrespective of sexual or mitotic origin (see *Materials and Methods*). As in linkage decay analysis, we considered the best-mapping chromosomes 1, 5, 21 and 26. Values of *ρ* for BM suggested ca. 3 meioses per 1,000 mitotic events in this group. By contrast, all approximations of *ρ* for EH and AR fell within confidence limits of the synthetic, non-recombinant *‘FSC_n’* control. These limits also contained *ρ* = 0 (Tbl. 2).

**Table 2.**
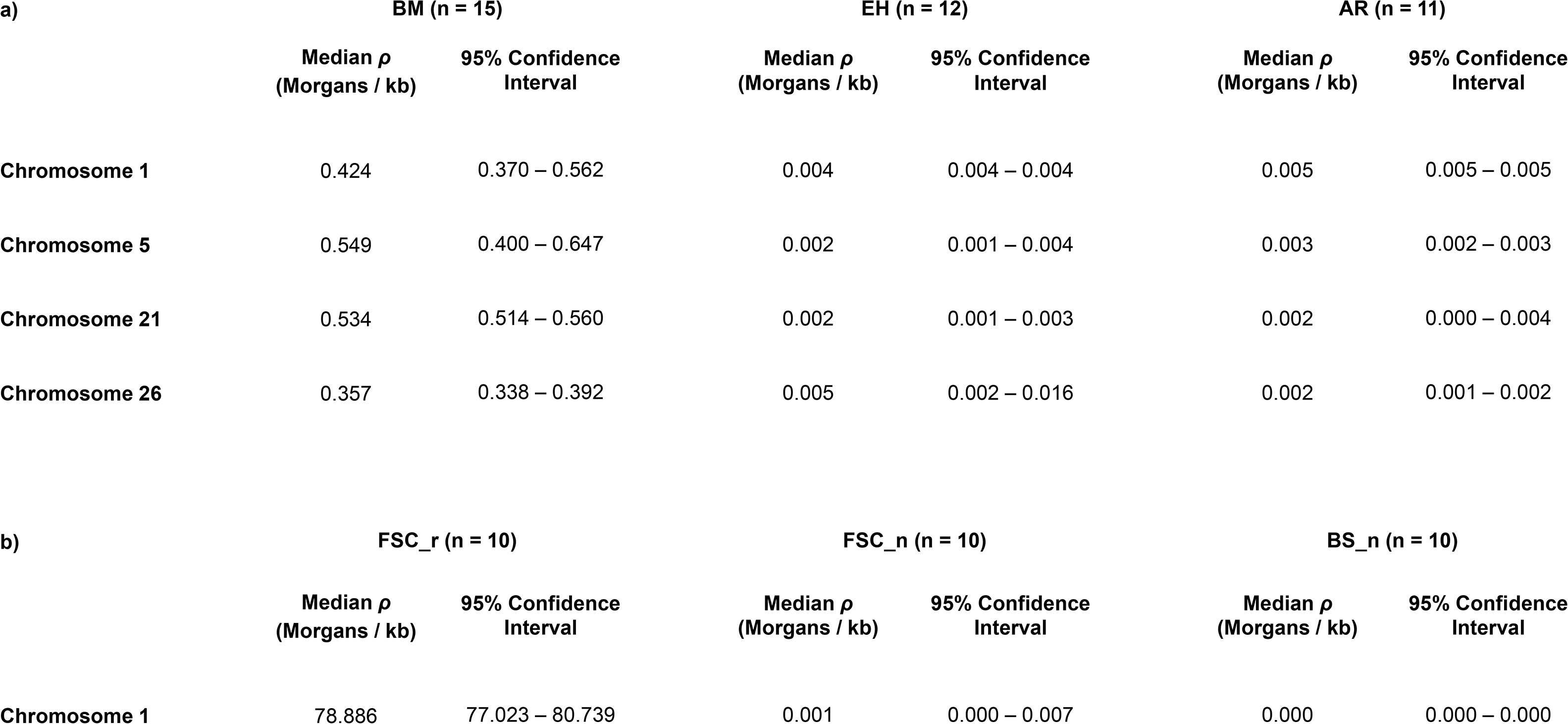
Composite-likelihood approximation of the population recombination parameter *ρ*, carried out using the ‘interval’ program in LDhat (89). (a) Results for *T. cruzi* I isolates from Bella Maria (BM), El Huayco (EH) and Ardanza (AR). (b) Results for simulated datasets with and without recombination (FSC_r: *ρ* = 128; FSC_n: *ρ* = 0; and BS_n: *ρ* = 0) demonstrate the sensitivity of the method.

### Evidence for chromosomal re-assortment, with or without linkage decay

Apart from disrupting sequence patterns within chromosomes, sexual mating breaks up associations between homologous chromosome-pairs. Given sufficient population diversity therefore, incongruent phylogenies built on the basis of chromosomal genotypes are expected. We came across many such cases in BM isolates of Cluster 1 (Fig. 3 and Fig. S7). Ancestries among several isolates from Cluster 2 also appeared incongruent at the chromosomal level. Discordant chromosomal phylogenies were conspicuous for EH samples THY_3973_CL3 and THY_4333_CL3, which clustered either with other samples from EH (e.g., for chromosome 1 and chromosome 5; Fig. 3) or with individuals from AR (e.g., see Fig. S7). GE samples TGM_4146_CL4 and TGM_4169_CL5 also varied notably in chromosome-by-chromosome phylogenetic comparison. These samples, which occupied peripheral positions in previous genome-wide analyses (e.g., NeighborNet results – Fig. 1a), generally paired with TCQ_3087 clones (e.g., for chromosome 1) but every so often cleaved to the base of the BM clade (e.g., for chromosome 5 and chromosome 19). We also recognized these varying affinities among EH, AR and GE isolates in supplementary DAPC trials at higher k-means solutions (e.g., see individual membership probabilities for k = 5 in Fig. S8).

**Figure 3.**
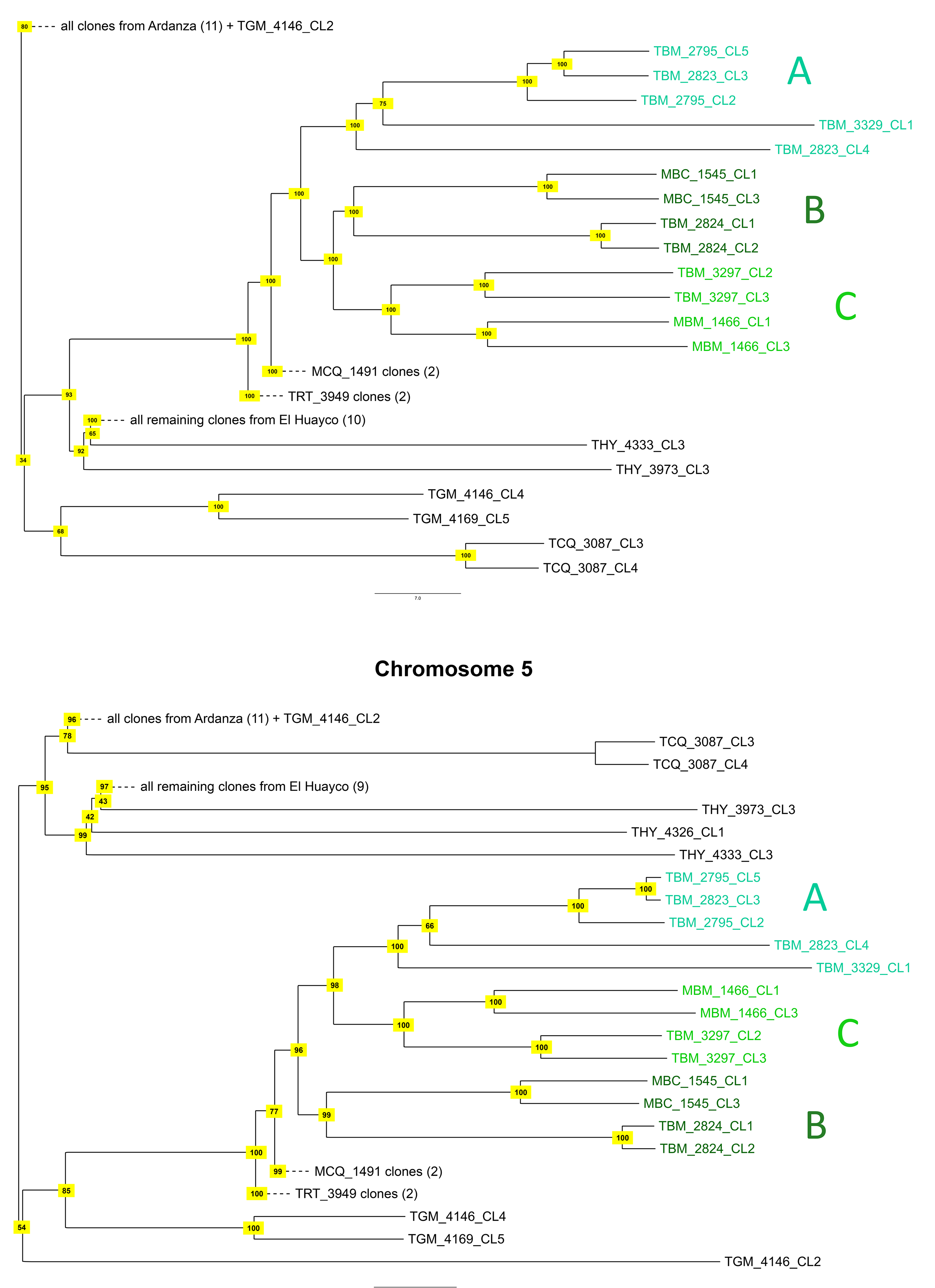
Incongruent neighbor-joining dendrograms exemplify independent chromosomal ancestry patterns among *T. cruzi* I isolates. Neighbor-joining results for chromosomes 1 and 5 discord at various well-supported nodes. For example, chromosome 1 presents a monophyletic clade composed of MBC_1545 + TBM_2824 (labelled ‘B’) and MBM_1466 + TBM_3297 clones (clade ‘C’) from the Bella Maria sampling region. An outgroup is formed by TBM_2795 + TBM_2823 + TBM_3329 clones (clade ‘A’), likewise from Bella Maria. On chromosome 5, clade A has changed places with clade C. The A+B monophylum occurs again elsewhere, e.g., on chromosome 19 (Fig. S7), while the B+C group makes appearances on, e.g., chromosomes 9 and 16. Nodes are labelled with support values from 100 bootstrap replicates. Dashed lines indicate collapsed clades.

### Indications of non-Mendelian hybridization

Of 80,052 sites that differed from the TcI-Sylvio reference in EH, 62,036 also differed in AR, and >50% of this mutual polymorphism occurred as fixed heterozygous SNPs across the two groups. These observations (as well as various congruences in prior descriptive metrics (e.g., see Tbl. 1)) provided first indications of shared ancestry across isolates of Cluster 2. Additional analyses of haplotype differentiation and somy structure furthered this premise with possible signs of non-Mendelian hybridization at the outset of highly heterozygous genomes found throughout Cluster 2.

In windowed haplotype analysis for Cluster 2, we found short patches of similarity in all between-group individual comparisons. Sudden drops in differentiation ranged within 60 - 180 kb and scattered unsystematically throughout the genome. These patterns of between-group ‘haplotype-sharing’ appeared similar to those within BM (see Fig. S4c-d (left panel) and Fig. S4b (right panel), respectively, for chromosome 1). Interestingly, patches of low differentiation between samples within groups of Cluster 2 often involved not one but two similar haplotypes, in other words, stretches of similar heterozygosity intervening otherwise dissimilar heterozygous sequence. Unlike haplotype-sharing described above, this ‘genotype-sharing’ within groups of Cluster 2 extended far beyond 180 kb, often to the end of the chromosome (e.g., see pairwise comparisons within EH for chromosome 1 in Fig. S4b (left panel)).

In somy analysis, chromosome-wide deviations in variant allele fraction and total read-depth suggested full-chromosome trisomies in ten samples from Cluster 2 (Fig. 4a-b), with highest rates in THY_3975, THY_4326 and THY_4332 clones (>10 trisomies each). Of 22 chromosomes with apparent trisomy, ten appeared trisomic in ≥5 samples, with similar biases apparent in EH and AR (e.g., chromosomes 19, 25 and 39). Regardless of ploidy status, chromosomes differed little from another in average heterozygosity at variable loci (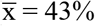 and σ = 4.6% in both EH and AR; Fig. S9), dropping below 30% only for chromosomes 14 and 41. We did not observe such rates of trisomy in Cluster 1 (BM). With the exception of TBM_2824 clones (trisomic for chromosomes 32 and 44), all BM isolates showed euploidy despite similar levels of intra-chromosomal read-depth variation. Most BM genomes showed severe reductions in sequencing coverage over chromosome 13. Such reductions did not occur in EH or AR (Fig. 4a). Somy plots for all samples are provided in Fig. S10. Fig. S10 includes results for the (euploid) reference strain as well as for all non-cloned samples, of which low-diversity infections such as MCQ_1491_MIX and THY_4327_MIX (see Fig. 1b) mirrored monosomy/trisomy deviations in correspondent clones.

**Figure 4.**
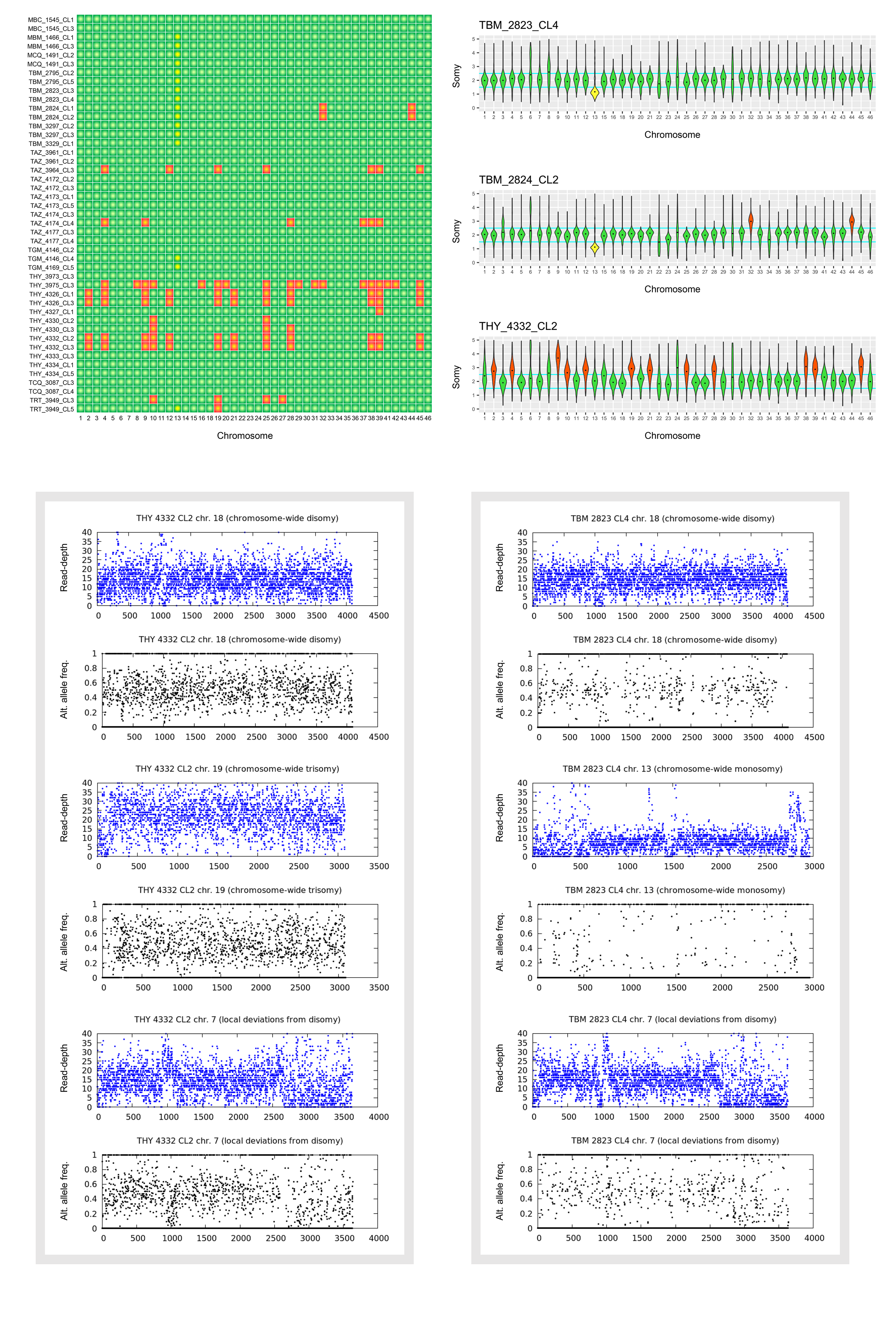
Population-level aneuploidy among *T. cruzi* I isolates. (**a**) We distinguished chromosomal and intra-chromosomal copy number variation by evaluating sequence read-depth kernel density distributions. For example, these distributions suggest multiple cases of whole-chromosome somy elevation (highlighted in red) for El Huayco isolate THY_4332_CL2 (bottom right). Several isolates from El Huayco and Ardanza present similar patterns (see Fig. S10 for more density plots), as summarized in the heatmap at left. Read-depth densities suggest few cases of whole-chromosome somy elevation for isolates from Bella Maria (e.g., TBM_2823_CL4 and TBM_2824_CL2 at right). However, mapping coverage drops dramatically (yellow) on chromosome 13 in most isolates of this group. (**b**) Chromosome-wide shifts in sequence read-depth (blue) and alternate allele frequency (black) support whole-chromosome aneuploidies inferred from read-depth kernel density distributions above. In El Huayco isolate THY_4332_CL2 (left column), for example, read-depth is elevated over the entirety of trisomic chromosome 19 (sequence positions are plotted on the x-axis). Alternate allele frequencies at heterozygous sites also distribute around values of 0.33 and 0.67 on this chromosome (as compared to frequencies around 0.50 on disomic chromosome 18). Cases of intra-chromosomal copy number variation for sample THY_4332_CL2 are marked by local shifts in read-depth and alternate allele frequency on chromosome 7. Comprehensive read-depth reduction on chromosome 13 is exemplified for Bella Maria isolate TBM_2823_CL4 (right column). Alternative allele frequency values of 0 (indicative of the reference allele) predominate on this chromosome. Patterns on chromosome 7 and 18 also point to intra-chromosomal copy number variation and stable disomy, respectively, for the TBM_2823_CL4 isolate.

### Mysterious homozygous migrants imply further parasexuality

TRT_3949 (sampled near EH but associated to Cluster 1) and TCQ_3087 (sampled in BM but associated to Cluster 2) were the only samples for which geographic and nuclear phylogenetic neighbors clearly did not match (Fig. 1, Fig. S1 and Tbl. S1). They also provided the dataset’s only cases of discordant nuclear vs. mitochondrial phylogenies. TRT_3949 clones carried a maxicircle genotype otherwise found only in Cluster 2 and TCQ_3087 clones carried a maxicircle genotype highly divergent to any other observed in the study (Fig. S11; see also mitochondrial read-depth aberration (nearly 3 standard deviations above average) for TCQ_3087 clones in Tbl. S3). These were the only samples for which homozygosity preponderated SNP-variation throughout the nuclear genome (Fig. 5a shows alternate allele frequency means in windowed analysis of chromosome 5; see analyses for other chromosomes in Fig. S3 and abundance of long runs of homozygosity in Tbl. S2). Making up just 10% total polymorphic loci, bi-allelic SNPs were found restricted to scattered patches, even the most heterozygous of which were predominated by homozygous sites. The extreme levels of overall homozygosity observed in these isolates did not attribute to certain chromosomes or to deviations in read-depth, and most homozygous sites (59 - 69%) matched TcI-Sylvio alleles.

**Figure 5.**
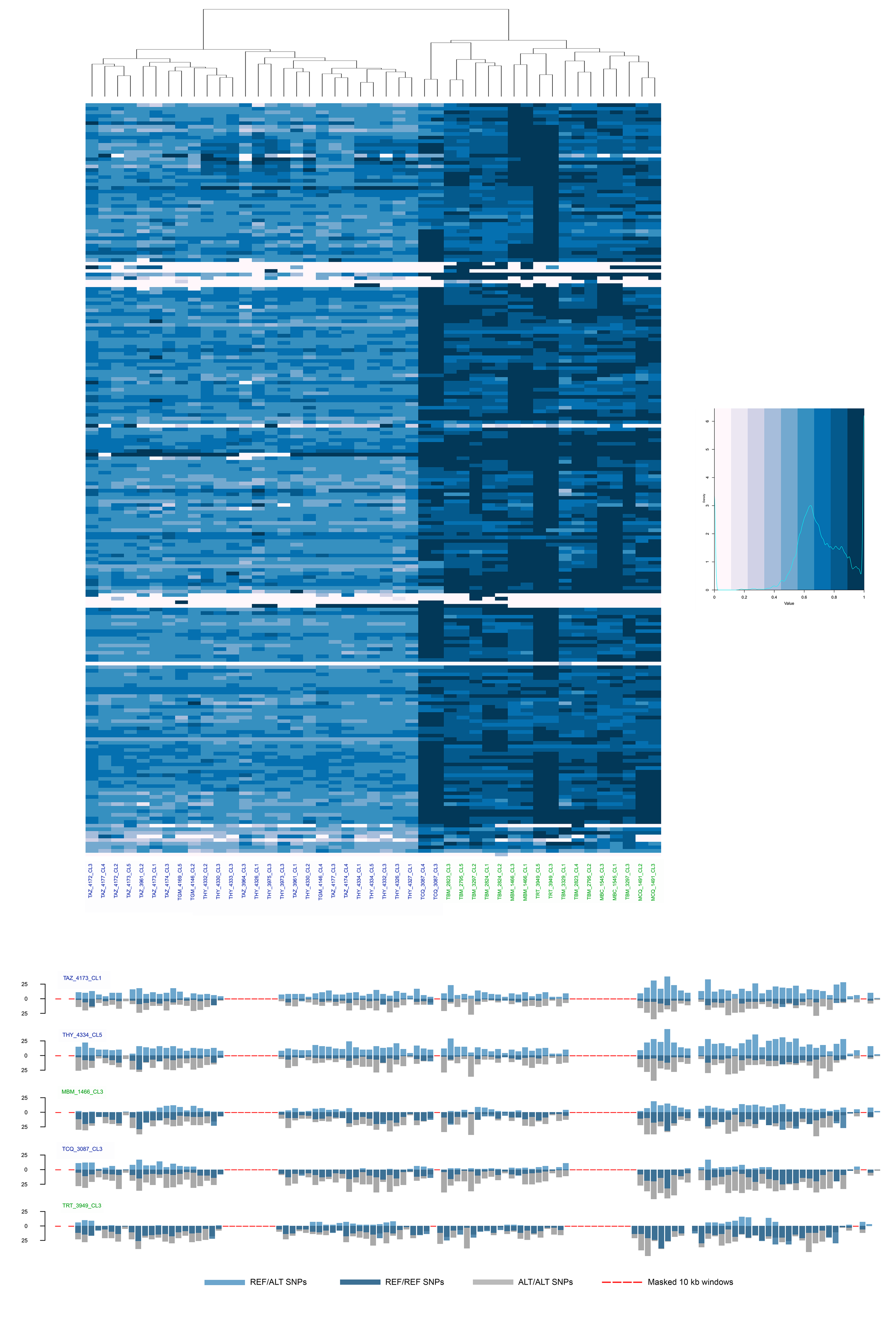
Genome-wide patterns of heterozygosity among *T. cruzi* I isolates. (**a**) Alternate allele frequency averages along chromosome 5 exemplify genome-wide patterns of heterozygosity in the dataset. Each column represents the chromosome of one *T. cruzi* I isolate. Green and blue labels at bottom signify Cluster 1 and 2 membership, respectively, as defined in Fig. 1. Rows within each column represent consecutive 5 kb sequence bins. Alternate allele frequency averages determine the color of each bin. Extreme homozygosity is apparent for TCQ_3087 and TRT_3949 clones. Isolates from Bella Maria tend to carry patchy heterozygosity while those of Cluster 2 appear highly heterozygous throughout the chromosome. Further examples are provided in Fig. S3. Gaps indicate regions without SNP sites. These are due in large part to masking (see below). (**b**) Heterozygous and homozygous SNP counts along chromosome 5 for various *T. cruzi* I isolates. Each bar quantifies SNP sites in a consecutive 10 kb window. Counts of heterozygous sites are colored in light blue and plotted upward on the y-axis. Counts of sites showing homozygosity for the reference allele or homozygosity for the alternate allele are plotted in the opposite direction and colored in dark blue and grey, respectively. Sites with additional alternate alleles are not counted. Red dashes indicate windows masked from analysis due to mapping uncertainty at >20% sites.

## DISCUSSION

### Meiotic sex and parasexual polyploidization in *T. cruzi*

Our comparative genomic analysis of 45 biologically cloned strains from an area of endemic transmission supports the remarkable conclusion that a) *T. cruzi* undergoes meiosis and b) that grossly disparate rates and mechanisms of genetic exchange occur simultaneously in a single *T. cruzi* focus.

In a subsection of the region (BM), signs of regular Mendelian sex abound. Genome-wide allele frequencies occur at HWE and ancestries among individuals fluctuate from chromosome to chromosome. Parasite genotypes on individual chromosomes appear equally mosaic: linkage decays to zero within 100 kb. We gauge that meiosis occurs more than once every 1000 reproductive events. In nearby populations EH, AR and GM, meiosis is essentially absent. Instead, the populations exhibit extreme heterozygosity and bear no signs of linkage decay. We do detect discordant chromosomal phylogenies among these parasites, but recombination estimates within chromosomes match those for synthetic, non-recombining controls.

Strikingly, several EH, AR and GM isolates also present extensive aneuploidy and long tracts of low differentiation, inconsistent with long-term clonality alone. When clonality represents the primary basis for genetic variation in a landscape, polymorphism among separate demes is expected to predominate in the form of random, private heterozygous sites – a result of long-term mutation accumulation, the so-called Meselson Effect (32). Ancestries are predicted to remain linked across non-homologous chromosomes (18) and discontinuities in genetic differentiation among individuals, if present, most likely result from mitotic gene conversion and thus manifest as absolute homozygosis on disomic chromosomes (33).

The highly heterozygous populations described herein do not fit this scenario. In light of disproportionate fixed heterozygosity observed among geographically separate demes, we instead consider full-genome hybridization as a primary driver of population genetic structure in EH, AR and GM. More specifically, we find that polyploid hybridization, closely linked to asexuality in various eukaryotes (34–37), better explains the karyotypic and genotypic patterns found in this study.

In the only genetic exchange event observed experimentally in *T. cruzi* to date (22), parental genomes fused to tetraploid hybrids and then began erosion back toward the disomic state. This ‘fusion-then-loss’ process resembles parasexual programs known from pathogenic fungi (*Candida* spp.) and allows for independent chromosomal ancestries without intra-chromosomal linkage decay (38, 39). Moreover, gene conversion in tetraploids can produce long tracts of increased identity on both homologues (i.e., referred to as ‘genotype-sharing’ in our results) without absolute homozygosis upon reduction to the disomic state (40). This is especially true when genome erosion is biased against the retention of similar homologues, a condition that aligns with our results (e.g., we observed elevations to average homozygosity in just two chromosomes, not in fifteen (33%) as would be expected in the case of random chromosome loss). Long-range genome conversion is thought to be particularly common in polyploid genomes (41) and often accompanies hybridization in *C. albicans*, for which selective chromosome retention also appears to occur (42, 43). While a number of other phenomena (e.g., failed meiosis (44), endo-reduplication (45)) apart from parasexual hybridization can comply with certain results from EH and AR (e.g., aneuploidy, independent chromosomal ancestry), they fail to reconcile others (e.g., absence of linkage decay, congruent aneuploidies) and include no empirically observed precedent.

### Rationale for reproductive polymorphism in *T. cruzi*

As an extraordinarily eclectic, stercocarian parasite, *T. cruzi* spreads very inefficiently between a great diversity of vectors and hosts. These also vary immensely in transmission competence (46, 47) and availability (48, 49) and occupy an array of disparate niches (including the domestic-sylvatic interface). The parasite’s life cycle thus likely represents a continuum of bottlenecks linked to frequent local extinction and recolonization events that increase levels of genetic drift and identity by descent (IBD). It may thus come to less surprise that observations of diffuse hybrid clonality around a restricted focus of sex in BM resemble spatio-temporal patterns of heterogony demonstrated in various other metapopulation systems (50–58). Facultative sex often coincides with strong metapopulation structure, in which sexual variants are predicted to occupy core habitat (where population subdivision and inbreeding depression is minimized) while asexual variants disperse more freely without fitness costs from close IBD during frequent founding events (59). In this study, mass elevation of Tajima’s *D* offers support for both hybridization and bottlenecked clonal propagation in generating an excess of intermediate-frequency variants over EH and AR. Genome-wide elevations of Tajima’s *D* observed at synonymous SNPs are expected when non-segregating, ancestral polymorphism outweighs the Meselson Effect (60, 61) – for instance, when clonal propagation from hybrid progenitors does not extend far into the past. Incomplete ploidy reduction (i.e., the presence of multiple trisomies) observed in several samples of this study also suggests hybridization as a relatively recent event.

### Low connectivity between Mendelian and non-Mendelian groups

Our genotype- and haplotype-based summaries of co-ancestry indicate that the meiotic parasite population in BM is genetically segregated from others with a more complex, non-Mendelian past that occur in nearby EH and AR. Genetic discontinuity occurs consistently for samples collected within a few kilometers distance and despite evidence for vector/host co-infection and migration between divergent groups. Putative migrants, or progeny thereof, appear highly homozygous with extensive (nuclear) homozygosity and, in the case of TCQ_3087 clones, exhibit extreme maxicircle sequence divergence and depth. Such observations are reminiscent of *L. major* crosses formed in non-native vectors (62)) and of irregular, biparental mitochondrial inheritance in transfected *T. b. brucei* (63).

Unexpected and poorly repeatable hybrid genomes have arisen on a number of occasions in experimental Tritryps research (20, 22, 64). Sensitivity to cryptic biochemical cues is clearly high, but the molecular signals that incite recombination and control mating compatibilities within these species remain essentially unknown (65). Our observations from the field do not identify such mechanisms but provide many relevant questions to explore. For instance, do ploidy barriers segregate non-meiotic cycles in *T. cruzi* as they do commonly in plants (66)? Is certain monosomy (e.g., recall chromosome 13 in BM isolates) associated with mating locus activation and sex? Similar mating cassette control is reported in *C. albicans* (67). Is extreme homozygosity a direct result of improper mating or a subsequent effect (gene conversion, selfing, etc.)? What are the adaptive processes that underpin switching between different reproductive modes?

## Conclusions

Our work now demonstrates irrefutable evidence of meiotic sex in *T. cruzi*, alongside complex parasexual reproduction and clonal parasite propagation. Complex mating structures are of acute relevance to Chagas disease control. Recombination implies that important epidemiological traits are transferable, not locked into stable subdivisions in space and time (for case in point, consider, e.g., SRA-gene transfer from T. *b. rhodesiense* to *T. b. brucei* (12, 13). Recombination also allows for pivotal disease features to arise *de novo* and is likely to have spurred major changes in *T. cruzi* transmission, including adaptation to the domestic niche (68, 69). Our data suggest that recombination may continue to transform contemporary disease cycles, as suggested for *Toxoplasma gondii* (70) and in *Leishmania* spp. (71–73). The proven presence of a sexual cycle in *T. cruzi* should now reinvigorate the hunt for the site of genetic exchange, as well as its cytological mechanism. An *in vitro* model for meiotic genetic exchange in *T. cruzi* will dramatically improve our ability to test the genetic basis of key epidemiologically relevant phenotypic traits – virulence, pathogenicity, drug resistance, etc. Determination of such traits may underpin future efforts to control and treat Chagas disease.

## MATERIALS AND METHODS

### Parasite collection and cloning

Trypanosomes were isolated from triatomines (*Rhodnius ecuadoriensis, Panstrongylus chinai, P. rufotuberculatus*, and *Triatoma carrioni*), rodents (*Rhipidomys leucodactylus*, *Simosciurus nebouxii*) and bats (*Artibeus fraterculus*) captured between 2011 and 2015 in eastern Loja Province, Ecuador. Capture coordinates, dates and ecotypes (i.e., domestic, peri-domestic or sylvatic) are provided in Tbl. S1 and associated protocols are detailed in previous studies led by the Center for Research on Health in Latin America (CISeAL) (74). Individual parasite cells were cloned on solid medium to derive single-strain colonies following Yeo et al. 2007 (75). Complementary to 19 non-cloned primary cultures, this process yielded 64 axenic monocultures for subsequent DNA extraction and sequencing.

### DNA sequencing and variant discovery

Genomic DNA was extracted from 83 *T. cruzi* cultures by isopropanol precipitation. DNA was sonicated and size-selected (median insert size = 198 nt; median absolute deviation = 69 nt) by covalent immobilization prior to paired-end sequencing on the Illumina HiSeq 2500 platform. To guide variant discovery from resultant 2 × 125 nt sequence reads, we optimized reference-mapping and SNP-calling pipelines using paired-end Illumina reads (kindly provided by Carlos Talavera-López) for *T. cruzi* TcI X10/1 (termed ‘TcI-Sylvio’ elsewhere in the text) against the newly available PacBio sequence for the same reference strain (29). Based on comparisons with TcI-Sylvio mapping results from various configurations in Smalt v.0.7.4 (we tested 12 - 14 kmer hash indexes and 2 - 8 base skip sizes), we chose to map samples using default settings (gap-open penalty = 6 and mismatch penalty = 4) in BWA-mem v0.7.3. We then sorted alignments with Samtools v0.1.18, marked PCR-duplicates with Picard v1.85 and identified single-nucleotide polymorphisms (SNPs) by local re-assembly with Genome Analysis Toolkit (GATK) v3.7.0 (76) (benchmarked for *L. donovani* in (77)). Individual records produced by the ‘HaplotypeCaller’ algorithm were subsequently merged for population-based genotype and likelihood assignment (GATK ‘GenotypeGVCFs’). Next, we calibrated variant filters by incrementally tightening thresholds for genotype quality (*QUAL*), read-depth (D) and local polymorphism density (C) until non-reference homozygous SNP-calls for TcI-Sylvio reached asymptotic decay. We then applied masks to exclude variant-calls where short-read alignment performed unreliably, that is, where ‘virtual mappability’ (*V*) (measured following Derrien et al. 2012 (78)) scored less than 100%. These regions represented areas of low sequence complexity and/or redundancy and made up large fractions of all reference chromosomes (Fig. S12a-b). With the above filters in place (*QUAL* > 1500; 10 > *D* < 100; *C* < 3 SNPs per 10 nt; *V* = 1), samples retained tens of thousands of homozygous alternative variant loci, whereas TcI-Sylvio Illumina vs. TcI-Sylvio PacBio showed just 58. Nevertheless, the guide-sample presented ca. 20,000 small insertions and ca. 1,000 small deletions relative to the reference. We placed an additional mask ±3 nt around these positions to avoid potential faults in the published genome. Final masking thus disqualified a total of 24 Mb (including all of chromosomes 17, 40 and 47) from polymorphism analysis. This highly conservative, diagnostic variant-screening approach also led us to exclude 24 low-depth samples for which genotypes could not be assigned at more than 40% variant sites. The final set of SNPs (in 59 samples) were annotated using the TcI-Sylvio annotation file (http://tritrypdb.org/common/downloads/Current_Release/TcruziSylvioX10-1/gff/data), and the effect of each variant on sequence expression was predicted using SnpEff v4.3t (79).

### Computational phasing of heterozygous SNP sites

Heterozygous SNP sites were phased over 30 iterations in BEAGLE v4.1 (80). The algorithm also imputes missing genotypes from identity-by-state segments found in the data (81). For haplotype co-ancestry and general phylogenetic analysis, we restricted imputation to sites containing information for >60% samples. Later, for quantitative phylogenetic analysis, however, we refrained from genotype imputation, i.e., used only sites with genotypes called in all individuals of the dataset.

### Detection of population genetic substructure

We used the Neighbor-Net algorithm in SplitsTree v4 (82) to visualize genome-wide phylogenetic relationships among samples in split network representation. Neighbor-Net extends Satou and Nei’s neighbor-joining algorithm to accommodate evolutionary processes such as recombination and hybridization that lead to non-treelike patterns of inheritance. We also optimized a general time-reversible (GTR) substitution model with ascertainment bias correction (for accurate branch lengths in the absence of constant sites) to construct phylogenies from proportions of non-shared alleles, i.e., considering two haplotypes per variant site. Haplotype concatenations were also used to derive a minimum-spanning network, the set of edges that links nodes (individuals) by the shortest possible cumulative distance (i.e., maximum-parsimony). We inferred genetic subdivisions in the sample-set by unsupervised k-means clustering and discriminant analysis of principle components (DAPC) (83). These analyses applied genetic distances as the proportion of non-shared genotypes at all variant loci (i.e., considering variants at the genotypic level), as did Neighbor-Net and subsequent measurements of *F_ST_*. After phasing heterozygous SNP sites (see above), we used fineSTRUCTURE v2.0.4 (31) to recover traces of identity-by-descent in similar haplotypes. This program was recently used to expose hybridization events in congeneric *T. congolense* (17), as well as to disentangle reticulate ancestries in the closely-related *L. donovani* complex (77). The built-in ‘Chromopainter’ algorithm (adapted from Li and Stephens (2003) (84)) constructs a semi-parametric summarization of co-ancestry among all pairs of individuals based on variable rates of haplotype-sharing and linkage disequilibrium across sample genomes. We applied fineSTRUCTURE over a uniform recombination map, running 6 · 10^5^ Markov chain Monte Carlo (MCMC) iterations (1 · 10^5^ iterations burn-in) and 4 · 10^5^ maximization steps in the final tree-building step. Following indications of mosaic inheritance in these analyses, we assessed phylogenetic (dis)continuity by comparing genotype-trees built for individual chromosomes as well as haplotype-trees built for phased segments (20 kb) within chromosomes. These analyses only considered windows with >200 SNP sites and brought general intra-chromosomal ancestry patterns into view, but we referred to separate distance matrices based on haplotypes phased without imputation (see previous section) to quantify changes in genetic similarity between windows within chromosomes. Both chromosomal and intra-chromosomal phylogenies were built using neighbor-joining as implemented in ‘ape’ v5.1 in R (85).

### Analyses of population genetic diversity and linkage

To assess population-level genetic diversity, we calculated site-wise nucleotide diversity (π), Watterson’s theta (*θ*) and *F_IS_* using ‘hierfstat’ v0.04-22 in R (86). *F_IS_* values rate heterozygosity observed within and between individuals, varying between −1 (all loci heterozygous for the same alleles) and 1 (all loci homozygous for different alleles). Values at 0 indicate Hardy-Weinberg equilibrium. We also measured rates of shared and private allele use (e.g., proportions of fixed heterozygous and singleton sites), assessed variant neutrality based on Tajima’s *D*, quantified haplotype diversity by counting unique haplotypes per 30 kb, and scanned for long runs of homozygosity using VCFtools v0.1.13 (87).

To determine linkage patterns within chromosomes 1, 5, 21, and 26 (the genome’s best-mappable chromosomes – see Fig. S12a-c), we recoded sample genotypes with values of 0, 1 or 2 to represent the number of non-reference alleles at each variant site. After filtering out all SNP-pairs separated by masked sequence (in effect, confining analysis to sites separated by <100 kb), we measured linkage (r^2^) as the correlation between genotypic allele counts in PLINK v1.90. We then binned r^2^ into distance classes (from 0 to 100 kb in increments of 2 kb) to visualize relationships between map distance and linkage disequilibrium in R. These analyses were also run separately on core sequence areas, as defined by areas of synteny among TcI-Sylvio, *T. b. brucei* and *L. major* annotated at http://tritrypdb.org. Intra-haplotypic recombination is unlikely to accompany meiotic crossover events in these areas of the genome (29).

### Estimation of meiotic vs. mitotic division

Following methods established to quantify complex microbial life cycles (88), we inferred the frequency of sex and clonality in *T. cruzi* isolates by comparing two different estimates of effective population size. The first estimate, *N_ρ_*, is based on recombinational diversity observed in the sample. *N_ρ_* represents the number of cells derived from mating, i.e., the number of zygotes present in the population, and is calculated as *ρ* / 4r (1 - *F*), where *ρ* denotes nucleotide covariation between sites, *r* denotes rate of recombination per bp per generation, and *F* represents Wright’s inbreeding coefficient. The second estimate, *N_θ_*, is based on mutational diversity observed in the sample. *N_θ_* represents the total population size, i.e., the number of cells irrespective of sexual or mitotic origin, and is calculated as *θ* (1 + *F*) / 4*μ*, where *θ* denotes nucleotide variation at single sites and *μ* denotes the rate of mutation per bp per generation. *N_ρ_* / *N_θ_* thus quantifies the frequency of meiotic reproduction in the population. To estimate this quotient from our sample, we derived *θ* from Watterson’s estimator at non-coding sites and derived *ρ* based on reversible-jump MCMC likelihood curves generated by the ‘interval’ program in LDhat (89). We used 1·10^7^ MCMC iterations with 2,000 updates between samples and block penalties set to five. We estimated *r* from the equation *r* = *0.043* · *S^−1.310^* and *μ* from the equation *μ* = *2.5866 η 10^10^ · S^0.584^*. These regression models were developed in Rogers et al. 2014 (19) based on the observation that genome size (*S*) correlates strongly to rates of recombination and mutation in unicellular eukaryotes. We validated *ρ* estimates by simulating input for LDhat in two ways. First, we created sequence alignment maps for ten non-recombinant individuals based on observed genotypes using BAMSurgeon v1.0.0 (90). Maps were set up for each individual by inserting fixed polymorphisms from the true sample set into TcI-Sylvio sequence reads, then spiking in random mutations at rates corresponding to the average number of pairwise differences in the observed data. Individual SNP records for the ten mutant alignment files were then compiled and merged to variant-call-format (VCF) in GATK as outlined above. In the second approach, we used fastsimcoal2 v5.2.8 (91) to simulate ten non-recombinant and ten recombinant genotypes, applying *r* and *μ* from above equations to an effective population of 100,000 diploid individuals under a finite-sites model of evolution for chromosome 1.

### Chromosomal somy analysis

To estimate somy levels for each sample, we first measured mean-read-depth for successive 1 kb windows spanning each chromosome. We then calculated the median of these windowed-depth-means (*m*), i.e., a ‘median-of-means’ (*M_m_*), for each chromosome. After testing at various distribution points, we let the 30^th^ percentile (*p30*) of (skewed) *M_m_* values represent expectations for the disomic state, estimating copy number for each chromosome by dividing its *M_m_* by the sample’s *p30* value and multiplying by two. This procedure produced estimates of disomy for all chromosomes of the TcI-Sylvio guide-sample and outperformed techniques based on different window-sizes as well as those refined according to sequence annotation (e.g., only single-copy genes) or mapping quality (data not shown). We validated cases of chromosomal copy number variation by plotting kernel densities for *m*, as well as by assessing raw depth and alternate allele frequencies across variant sites. True, whole-chromosomal trisomy, for example, should translate to chromosome-wide elevations in read-depth and reductions in minor allele contributions to ca. 33% (i.e., one ‘B’ and two ‘A’ alleles – and, in cases of tri-allelism, one of each ‘A’, ‘B’ and ‘C’ alleles) at all heterozygous (i.e., ‘A/B/B’ or ‘A/B/C’) sites. Intra-chromosomal amplification, in contrast, should create local shifts in read-depth and allelic composition within chromosomes.

## ACKNOWLEDGEMENTS

This study was funded by the Division of Microbiology and Infectious Diseases, the National Institute of Allergy and Infectious Diseases and the National Institutes of Health (DMID/NIADID/NIH grants AI077896-01 and AI105749-01A1); the NIH-Fogarty Global Infectious Disease Training Program (grant TW008261); the Pontifical Catholic University of Ecuador (I13048, J13033, K13063 and L13225); and the Scottish Universities Life Science Alliance (SULSA). The funders had no role in study design, data collection and analysis, decision to publish, or preparation of the manuscript.

---

**Figure S1. Additional representations of phylogenomic relationships among *T. cruzi* I isolates.** (**a**) Data are represented as a minimum-spanning network (92). Multi-furcating nodes are arranged such that cumulative edge distance is minimized among samples. Pairwise genetic distances are haplotype-based, defined as the proportion of non-shared alleles across all SNP sites for which genotypes are called for all individuals.

(**b**) Maximum-likelihood phylogeny for all *T. cruzi* I isolates under a general time-reversible (GTR) substitution model with ascertainment bias. Pairwise distances are also haplotype-based as above.

**Figure S2. Rates of individual homozygosity relative to Hardy-Weinberg expectations in *T. cruzi* I populations.**

Genome-wide density distributions of Wright’s inbreeding coefficient *F_IS_* are plotted for *T. cruzi* I isolates from Bella Maria, El Huayco and Ardanza. Mean and standard deviation are also given for each group.

**Figure S3. Patterns of heterozygosity on chromosomes 9 and 21 for *T. cruzi* I isolates.**

**(a-b)** Alternate allele frequency averages along chromosome 9 and 21, respectively, are plotted as in Fig. 5a.

**Figure S4. Patterns of haplotype differentiation within chromosomes of *T. cruzi* I isolates.**

(**a**) Number of distinct haplotypes found in phased sequence alignment at window sizes increasing from 0 to 100 kb. Boxplots represent isolates from Bella Maria (green), El Huayco (dark blue) and Ardanza (light blue).

(**b**) Within-group pairwise haplotype differentiation of *T. cruzi* I isolates from El Huayco (left) and Bella Maria (right). Each row (i.e., biplot) represents the comparison of a reference individual (e.g., THY_4334_CL5, chosen to represent El Huayco based on phylogenetic positions in Fig. 1a and Fig. S1) to another individual specified at right. Light green bars indicate pairwise dissimilarities in consecutive 60 kb sequence windows along phased haplotype A. Opposite bars (dark green) quantify pairwise dissimilarities for phased haplotype B. Grey color indicates regions of low polymorphism (<20 SNP sites per 60 kb), due in part to masking and thus excluded from analysis. Pairwise dissimilarities are given as log-scaled Euclidean distances.

**(c-d)** Haplotype differentiation of *T. cruzi* I isolates from El Huayco (left) and Bella Maria (right) relative to isolates from other groups. These between-group pairwise comparisons are presented in the same way as within-group pairwise comparisons are presented above.

**Figure S5. Rates of nucleotide diversity relative to neutral expectations in *T. cruzi* I populations.**

Histograms of Tajima’s *D* for *T. cruzi* I isolates from Bella Maria, El Huayco and Ardanza. Values are calculated from 50 kb sequence bins throughout the genome.

**Figure S6. Linkage decay in core sequence regions of chromosome 1 in *T. cruzi* I isolates from the Bella Maria sampling region.**

Core sequence regions are defined as areas of synteny among TcI-Sylvio, *T. b. brucei* and *L. major* reference genomes. Only short map distance classes (0 - 1.2 kb) are considered due to low sample size in this restricted analysis. Presentation is otherwise analogous to that in Fig. 2a.

**Figure S7. Neighbor-joining relationships on chromosome 19 among *T. cruzi* I isolates.**

Red arrows suggest areas for comparison to Fig. 3, which presents relationships on chromosomes 1 and 5 in analogous format.

**Figure S8. Nonparametric population clustering of *T. cruzi* I isolates.**

(**a**) Bayesian Information Criterion (BIC) scores of different k-means clustering solutions for population assignment of *T. cruzi* I isolates. Ward’s criterion (30) was applied for objective selection of k.

(**b**) Discriminant analysis of principle components (DAPC) membership probabilities for k = 2, k = 5 and k = 6. Latter k-means solutions allow for additional partitioning of sample genetic diversity but do not necessarily represent true population subdivision.

**Figure S9. Average heterozygosity (per chromosome) for *T. cruzi* I isolates from El Huayco and Ardanza sampling regions.**

Average heterozygosity values fall between 40 and 50% for most chromosomes. Only chromosomes 14 and 41 show substantial increases in homozygosity.

**Figure S10. Somy estimates for cloned and non-cloned *T. cruzi* I samples.**

Somy variation among samples was inferred from sequence read-depth kernel density distributions. Results for the TcI-Sylvio reference validate the procedure. Results for non-cloned, low-diversity infections (e.g., MCQ_1491_MIX and THY_4327_MIX) indicate that aneuploidies in cloned samples are not a consequence of plate-cloning procedures in the lab.

**Figure S11. Mitochondrial phylogenies for *T. cruzi* I isolates.**

(**a**) Maxicircle sequence variation among all samples except TAZ_4172_CL3 (missing information at 44% sites) is represented as a TCS network (93). Black tick marks between nodes indicate number of mutations between different maxicircle genotypes. Node sizes correspond to the number of samples represented by the particular maxicircle variant. Green and blue fill colors signify Cluster 1 and 2 membership, respectively, as defined from nuclear phylogenetic analysis (Fig. 1). TCQ_3087 clones are extremely anomalous with 668 diagnostic SNPs relative to other isolates of Cluster 2.

(**b**) Complementary neighbor-joining analyses of cytochrome b sequences do not readily attribute extreme maxicircle divergence in TCQ_3087 clones to kinetoplast inheritance from non-TcI DTUs. Additional sequences applied in the dendrogram are detailed in (94) and (95).

**Figure S12. TcI-Sylvio reference evaluation and masking.**

**(a)** Masks applied to the TcI-Sylvio reference genome based primarily on ‘virtual mappability’ (78). Final masking disqualified a total of 24 Mb (including all of chromosomes 17, 40 and 47) of 42 Mb from polymorphism analysis.

**(b-c)** Proportions of ‘mappable’ (78), ‘unique’ (determined by self-blasting) and gap content on TcI-Sylvio reference chromosomes are indicated in light grey, dark grey and blue, respectively. Red bars distinguish chromosomes excluded from analysis based on these metrics.

## REFERENCES

1. Pérez-Molina JA, Molina I (2018) Chagas disease. The Lancet 391(10115):82–94.

2. Coura JR, Dias JCP (2009) Epidemiology, control and surveillance of Chagas disease: 100 years after its discovery. Memórias do Instituto Oswaldo Cruz 104:31–40.

3. Rassi A Jr, Rassi A, Marin-Neto JA (2010) Chagas disease. Lancet 375(9723): 1388–1402.

4. Bern C, Martin DL, Gilman RH (2011) Acute and congenital Chagas disease. Adv Parasitol 75:19–47.

5. Monteiro FA, Weirauch C, Felix M, Lazoski C, Abad-Franch F (2018) Chapter Five - Evolution, Systematics, and Biogeography of the Triatominae, Vectors of Chagas Disease. Advances in Parasitology, eds Rollinson D, Stothard JR (Academic Press), pp 265–344.

6. Yeo M, et al. (2005) Origins of Chagas disease: Didelphis species are natural hosts of Trypanosoma cruzi I and armadillos hosts of Trypanosoma cruzi II, including hybrids. International Journal for Parasitology 35(2):225–233.

7. Simpson AGB, Stevens JR, Lukeš J (2006) The evolution and diversity of kinetoplastid flagellates. Trends in Parasitology 22(4): 168–174.

8. Lukes J, et al. (2002) Kinetoplast DNA network: evolution of an improbable structure. Eukaryotic Cell 1(4):495–502.

9. Clayton C (2013) The Regulation of Trypanosome Gene Expression by RNA-Binding Proteins. PLOSPathogens 9(11):e1003680.

10. Ramesh MA, Malik S-B, Logsdon JM (2005) A Phylogenomic Inventory of Meiotic Genes: Evidence for Sex in Giardia and an Early Eukaryotic Origin of Meiosis. Current Biology 15(2):185–191.

11. Ramírez JD, Llewellyn MS (2014) Reproductive clonality in protozoan pathogens—truth or artefact? Mol Ecol 23(17):4195–4202.

12. Balmer O, Beadell JS, Gibson W, Caccone A (2011) Phylogeography and Taxonomy of Trypanosoma brucei. PLOS Neglected Tropical Diseases 5(2):e961.

13. Gibson W (2015) Liaisons dangereuses: sexual recombination among pathogenic trypanosomes. Res Microbiol 166(6):459–466.

14. Tait A (1980) Evidence for diploidy and mating in trypanosomes. Nature 287(5782):536–538.

15. MacLeod A, et al. (2005) Allelic segregation and independent assortment in T. brucei crosses: proof that the genetic system is Mendelian and involves meiosis. Mol Biochem Parasitol 143(1):12–19.

16. Peacock L, Bailey M, Carrington M, Gibson W (2014) Meiosis and Haploid Gametes in the Pathogen Trypanosoma brucei. Current Biology 24(2):181–186.

17. Tihon E, Imamura H, Dujardin J-C, Abbeele JVD, Broeck FV den (2017) Discovery and genomic analyses of hybridization between divergent lineages of Trypanosoma congolense, causative agent of Animal African Trypanosomiasis. Molecular Ecology 26(23):6524–6538.

18. Weir W, et al. (2016) Population genomics reveals the origin and asexual evolution of human infective trypanosomes. eLife Sciences 5:e11473.

19. Rogers MB, et al. (2014) Genomic Confirmation of Hybridisation and Recent Inbreeding in a Vector-Isolated Leishmania Population. PLOS Genetics 10(1):e1004092.

20. Akopyants NS, et al. (2009) Demonstration of genetic exchange during cyclical development of Leishmania in the sand fly vector. Science 324(5924):265–268.

21. Messenger LA, Miles MA (2015) Evidence and importance of genetic exchange among field populations of Trypanosoma cruzi. Acta Trop 151:150–155.

22. Gaunt MW, et al. (2003) Mechanism of genetic exchange in American trypanosomes. Nature 421(6926):936–939.

23. Lewis MD, et al. (2009) Flow cytometric analysis and microsatellite genotyping reveal extensive DNA content variation in Trypanosoma cruzi populations and expose contrasts between natural and experimental hybrids. Int J Parasitol 39(12):1305–1317.

24. Llewellyn MS, et al. (2011) Extraordinary Trypanosoma cruzi diversity within single mammalian reservoir hosts implies a mechanism of diversifying selection. Int J Parasitol 41(6–10):609–614.

25. Minning TA, Weatherly DB, Flibotte S, Tarleton RL (2011) Widespread, focal copy number variations (CNV) and whole chromosome aneuploidies in Trypanosoma cruzi strains revealed by array comparative genomic hybridization. BMC Genomics 12:139.

26. Souza RT, et al. (2011) Genome Size, Karyotype Polymorphism and Chromosomal Evolution in Trypanosoma cruzi. PLOS ONE 6(8):e23042.

27. Tibayrenc M, Kjellberg F, Ayala FJ (1990) A clonal theory of parasitic protozoa: the population structures of Entamoeba, Giardia, Leishmania, Naegleria, Plasmodium, Trichomonas, and Trypanosoma and their medical and taxonomical consequences. Proc Natl Acad Sci U S A 87(7):2414–2418.

28. Tibayrenc M, Ayala FJ (2017) Trypanosoma cruzi and the model of predominant clonal evolution. American Trypanosomiasis Chagas Disease (Second Edition) (Elsevier, London), pp 475–495.

29. Talavera-Lopez C, et al. (2018) Repeat-driven generation of antigenic diversity in a major human pathogen, Trypanosoma cruzi. bioRxiv:283531.

30. Ward JH Jr (1963) Hierarchical Grouping to Optimize an Objective Function. Journal of the American Statistical Association 58(301):236.

31. Lawson DJ, Hellenthal G, Myers S, Falush D (2012) Inference of Population Structure using Dense Haplotype Data. PLOS Genetics 8(1):e1002453.

32. Welch DBM, Meselson M (2000) Evidence for the Evolution of Bdelloid Rotifers Without Sexual Reproduction or Genetic Exchange. Science 288(5469): 1211–1215.

33. LaFave MC, Sekelsky J (2009) Mitotic Recombination: Why? When? How? Where? PLoS Genet 5(3). doi: 10.1371/journal.pgen.1000411.

34. Ament-Velásquez SL, et al. (2016) Population genomics of sexual and asexual lineages in fissiparous ribbon worms (Lineus, Nemertea): hybridization, polyploidy and the Meselson effect. Molecular Ecology 25(14):3356–3369.

35. Mogie M (1986) On the Relationship Between Asexual Reproduction and Polyploidy. J Theor Biol 122(4):493–498.

36. Lampert KP, Schartl M (2008) The origin and evolution of a unisexual hybrid: Poecilia formosa. Philos Trans R Soc Lond, B, Biol Sci 363(1505):2901–2909.

37. Lovell JT, et al. (2013) On the origin and evolution of apomixis in Boechera. Plant Reprod 26(4):309–315.

38. Bennett RJ, Johnson AD (2003) Completion of a parasexual cycle in Candida albicans by induced chromosome loss in tetraploid strains. The EMBO Journal 22(10):2505–2515.

39. Nieuwenhuis BPS, James TY (2016) The frequency of sex in fungi. Philos Trans R Soc Lond B Biol Sci 371(1706). doi:10.1098/rstb.2015.0540.

40. Bennett RJ, Forche A, Berman J (2014) Rapid Mechanisms for Generating Genome Diversity: Whole Ploidy Shifts, Aneuploidy, and Loss of Heterozygosity. Cold Spring Harb Perspect Med 4(10):a019604.

41. Sémon M, Wolfe KH (2007) Consequences of genome duplication. Curr Opin Genet Dev 17(6):505–512.

42. Forche A, et al. (2008) The Parasexual Cycle in Candida albicans Provides an Alternative Pathway to Meiosis for the Formation of Recombinant Strains. PLOS Biology 6(5):e110.

43. Bennett RJ (2015) The Parasexual Lifestyle of Candida albicans. Curr Opin Microbiol 28:10–17.

44. Kavanaugh LA, Fraser JA, Dietrich FS (2006) Recent Evolution of the Human Pathogen Cryptococcus neoformans by Intervarietal Transfer of a 14-Gene Fragment. Mol Biol Evol 23(10):1879–1890.

45. Lee HO, Davidson JM, Duronio RJ (2009) Endoreplication: polyploidy with purpose. Genes Dev 23(21):2461–2477.

46. Zeledón R, Alvarado R, Jirón LF (1977) Observations on the feeding and defecation patterns of three triatomine species (Hemiptera: Reduviidae). Acta Trop 34(1):65–77.

47. Gürtler RE, Cardinal MV (2015) Reservoir host competence and the role of domestic and commensal hosts in the transmission of Trypanosoma cruzi. Acta Trop 151:32–50.

48. Levy MZ, et al. (2015) Bottlenecks in domestic animal populations can facilitate the emergence of Trypanosoma cruzi, the aetiological agent of Chagas disease. Proc R Soc B 282(1810):20142807.

49. Grijalva MJ, Terán D, Dangles O (2014) Dynamics of Sylvatic Chagas Disease Vectors in Coastal Ecuador Is Driven by Changes in Land Cover. PLOS Neglected Tropical Diseases 8(6):e2960.

50. Li X-W, Wang P, Fail J, Shelton AM (2015) Detection of Gene Flow from Sexual to Asexual Lineages in Thrips tabaci (Thysanoptera: Thripidae). PLOS ONE 10(9):e0138353.

51. Sandrock C, Schirrmeister BE, Vorburger C (2011) Evolution of reproductive mode variation and host associations in a sexual-asexual complex of aphid parasitoids. BMC Evolutionary Biology 11:348.

52. Verduijn MH, Dijk PJV, Damme JMMV Distribution, phenology and demography of sympatric sexual and asexual dandelions (Taraxacum officinale s.l.): geographic parthenogenesis on a small scale. Biological Journal of the Linnean Society 82(2):205–218.

53. Vorburger C, Lancaster M, Sunnucks P (2003) Environmentally related patterns of reproductive modes in the aphid Myzus persicae and the predominance of two “superclones” in Victoria, Australia. Mol Ecol 12(12):3493–3504.

54. Lehto MP, Haag CR (2010) Ecological differentiation between coexisting sexual and asexual strains of Daphnia pulex. J Anim Ecol 79(6): 1241–1250.

55. Booij CJH, Guldemond JA (1984) Distributional and Ecological Differentiation Between Asexual Gynogenetic Planthoppers and Related Sexual Species of the Genus muellerianella (Homoptera, Delphacidae). Evolution 38(1):163–175.

56. Amat I, Castelo M, Desouhant E, Bernstein C (2006) The influence of temperature and host availability on the host exploitation strategies of sexual and asexual parasitic wasps of the same species. Oecologia 148(1):153–161.

57. Halkett F, Kindlmann P, Plantegenest M, Sunnucks P, Simon JC (2006) Temporal differentiation and spatial coexistence of sexual and facultative asexual lineages of an aphid species at mating sites. J Evol Biol 19(3):809–815.

58. Schenck RA, Vrijenhoek RC (1986) Spatial and temporal factors affecting coexistence among sexual and clonal forms of Poeciliopsis. Evolution 40(5): 1060–1070.

59. Haag CR, Ebert D (2004) A new hypothesis to explain geographic parthenogenesis. Annales Zoologici Fennici 41(4):539–544.

60. Tajima F (1989) Statistical method for testing the neutral mutation hypothesis by DNA polymorphism. Genetics 123(3):585–595.

61. Wakeley J, Aliacar N (2001) Gene Genealogies in a Metapopulation. Genetics 159(2):893–905.

62. Inbar E, et al. (2013) The Mating Competence of Geographically Diverse Leishmania major Strains in Their Natural and Unnatural Sand Fly Vectors. PLOS Genetics 9(7):e1003672.

63. Gibson W, Peacock L, Ferris V, Williams K, Bailey M (2008) The use of yellow fluorescent hybrids to indicate mating in Trypanosoma brucei. Parasites & Vectors 1:4.

64. Peacock L, Ferris V, Bailey M, Gibson W (2008) Fly transmission and mating of Trypanosoma brucei strain 427. Molecular and Biochemical Parasitology 160(2): 100–106.

65. Peacock L, Ferris V, Bailey M, Gibson W (2014) Mating compatibility in the parasitic protist Trypanosoma brucei. Parasit Vectors 7:78.

66. Crawford DJ (2004) The Role of Chromosomal Change in Plant Evolution. Oxford Series in Ecology and Evolution. By Donald A Levin. The Quarterly Review of Biology 79(3):311–312.

67. Zhang N, et al. (2015) Selective Advantages of a Parasexual Cycle for the Yeast Candida albicans. Genetics 200(4): 1117–1132.

68. Machado CA, Ayala FJ (2001) Nucleotide sequences provide evidence of genetic exchange among distantly related lineages of Trypanosoma cruzi. Proc Natl Acad Sci USA 98(13):7396–7401.

69. Lewis MD, et al. (2011) Recent, Independent and Anthropogenic Origins of Trypanosoma cruzi Hybrids. PLOS Neglected Tropical Diseases 5(10):e1363.

70. Grigg ME, Bonnefoy S, Hehl AB, Suzuki Y, Boothroyd JC (2001) Success and virulence in Toxoplasma as the result of sexual recombination between two distinct ancestries. Science 294(5540): 161–165.

71. Ravel C, et al. (2006) First report of genetic hybrids between two very divergent Leishmania species: Leishmania infantum and Leishmania major. International Journal for Parasitology 36(13):1383–1388.

72. Nolder D, Roncal N, Davies CR, Llanos-Cuentas A, Miles MA (2007) Multiple hybrid genotypes of Leishmania (viannia) in a focus of mucocutaneous Leishmaniasis. Am J Trop Med Hyg 76(3):573–578.

73. Volf P, et al. (2007) Increased transmission potential of Leishmania major/Leishmania infantum hybrids. International Journal for Parasitology 37(6):589–593.

74. Grijalva MJ, et al. (2015) Comprehensive Survey of Domiciliary Triatomine Species Capable of Transmitting Chagas Disease in Southern Ecuador. PLOS Neglected Tropical Diseases 9(10):e0004142.

75. Yeo M, et al. (2007) Resolution of multiclonal infections of Trypanosoma cruzi from naturally infected triatomine bugs and from experimentally infected mice by direct plating on a sensitive solid medium. Int J Parasitol 37(1): 111–120.

76. DePristo MA, et al. (2011) A framework for variation discovery and genotyping using next-generation DNA sequencing data. Nat Genet 43(5):491–498.

77. Imamura H, et al. (2016) Evolutionary genomics of epidemic visceral leishmaniasis in the Indian subcontinent. eLife Sciences 5:e12613.

78. Derrien T, et al. (2012) Fast Computation and Applications of Genome Mappability. PLoS One 7(1). doi: 10.1371/journal.pone.0030377.

79. Cingolani P, et al. (2012) A program for annotating and predicting the effects of single nucleotide polymorphisms, SnpEff: SNPs in the genome of Drosophila melanogaster strain w1118; iso-2; iso-3. Fly (Austin) 6(2):80–92.

80. Browning SR, Browning BL (2007) Rapid and Accurate Haplotype Phasing and Missing-Data Inference for Whole-Genome Association Studies By Use of Localized Haplotype Clustering. The American Journal of Human Genetics 81(5):1084–1097.

81. Browning BL, Browning SR (2016) Genotype Imputation with Millions of Reference Samples. The American Journal of Human Genetics 98(1): 116–126.

82. Huson DH, Bryant D (2006) Application of phylogenetic networks in evolutionary studies. Mol Biol Evol 23(2):254–267.

83. Jombart T, Devillard S, Balloux F (2010) Discriminant analysis of principal components: a new method for the analysis of genetically structured populations. BMC Genetics 11:94.

84. Li N, Stephens M (2003) Modeling linkage disequilibrium and identifying recombination hotspots using single-nucleotide polymorphism data. Genetics 165(4):2213–2233.

85. Paradis E, Claude J, Strimmer K (2004) APE: Analyses of Phylogenetics and Evolution in R language. Bioinformatics 20(2):289–290.

86. Goudet J (2005) hierfstat, a package for r to compute and test hierarchical F-statistics. Molecular Ecology Notes 5(1):184–186.

87. Danecek P, et al. (2011) The variant call format and VCFtools. Bioinformatics 27(15):2156–2158.

88. Tsai IJ, Bensasson D, Burt A, Koufopanou V (2008) Population genomics of the wild yeast Saccharomyces paradoxus: Quantifying the life cycle. PNAS 105(12):4957–4962.

89. McVean GAT, et al. (2004) The fine-scale structure of recombination rate variation in the human genome. Science 304(5670):581–584.

90. Ewing AD, et al. (2015) Combining tumor genome simulation with crowdsourcing to benchmark somatic single-nucleotide-variant detection. Nat Methods 12(7):623–630.

91. Excoffier L, Dupanloup I, Huerta-Sánchez E, Sousa VC, Foll M (2013) Robust demographic inference from genomic and SNP data. PLoS Genet 9(10):e1003905.

92. Excoffier L, Smouse PE (1994) Using Allele Frequencies and Geographic Subdivision to Reconstruct Gene Trees within a Species: Molecular Variance Parsimony. Genetics 136(1):343–359.

93. Clement M, Posada D, Crandall KA (2000) TCS: a computer program to estimate gene genealogies. Mol Ecol 9(10): 1657–1659.

94. Messenger LA, et al. (2012) Multiple mitochondrial introgression events and heteroplasmy in trypanosoma cruzi revealed by maxicircle MLST and next generation sequencing. PLoS Negl Trop Dis 6(4):e1584.

95. Westenberger SJ, et al. (2006) Trypanosoma cruzi mitochondrial maxicircles display species- and strain-specific variation and a conserved element in the non-coding region. BMC Genomics 7:60.

